# Single-nuclei transcriptomics of dog hippocampus reveals the distinct cellular mechanism of domestication

**DOI:** 10.1101/2022.01.29.478120

**Authors:** Qi-Jun Zhou, Xingyan Liu, Longlong Zhang, Rong Wang, Tingting Yin, Xiaolu Li, Guimei Li, Yuqi He, Zhaoli Ding, Pengcheng Ma, Shi-Zhi Wang, Bingyu Mao, Shihua Zhang, Guo-Dong Wang

## Abstract

The process of dog domestication leads to dramatic differences in behavioral traits compared to grey wolves. A class of putative positively selected genes is related to learning and memory, for instance, long-term potentiation and long-term depression. In this study, we constructed a single-nuclei transcriptomic atlas of the dog hippocampus to illustrate its cell types, cell lineage, and molecular features. Using the transcriptomes of 105,057 single-nuclei from the hippocampus of a Beagle dog, we identified 26 cell clusters and a putative trajectory of oligodendrocyte development. Comparative analysis revealed a significant convergence between dog differentially expressed genes (DEGs) and putative positively selected genes (PSGs). 40 putative PSGs were DEGs in the glutamatergic neurons, especially in the cluster 14, which is related to the regulation of nervous system development. In summary, this study provided a blueprint to understand the cellular mechanism of dog domestication.

## Introduction

The process of animal domestication leads to dramatic differences in behavioral and morphological traits compared to their wild ancestors (Plassais, et al. 2019; Wang, et al. 2020). Shared with the human being for parallel evolution and convergence evolution in the history of early hunting-gathering time and the recent living environment change from agrarian societies to modern urban lifestyle (Wang, et al. 2013; Wang, et al. 2016; Cao, et al. 2021; Liu, et al. 2021), domestic dogs have been undergoing strong artificial selection and resulted in approximately 450 globally recognized breeds, which makes them the most variable mammalian species on Earth (Hajeski 2016). Genome-wide scans for positive selection revealed that the behavioral and neurological traits are likely changed subsequently with the process of domestication. For instance, putative positively selected genes (PSGs) in dogs were reported linked to neural crest and central nervous system development (Pendleton, et al. 2018). The gene expression in brains showed a strong difference of putative PSGs in neurons on learning, memory, and behavior between domestic animals and their wild relatives (Li, et al. 2013; Li, et al. 2014).

The hippocampus is an important part of the limbic system in the brain that has an essential role in the consolidation of memory (Nadel and Moscovitch 1997; Bird and Burgess 2008). A class of domesticated genes is related to hippocampus synaptic long-term potentiation and long-term depression (Wang, et al. 2016). For instance, *DKKL1* is expressed in the ventral hippocampus and is associated with increased susceptibility to social defeat stress in mice (Bagot, et al. 2016). The single-cell RNA sequencing (scRNA-seq) has been used to identify the cell type atlas (Zeisel, et al. 2015), revealed cellular and molecular dynamics in neurogenesis (Artegiani, et al. 2017; Hochgerner, et al. 2018), and decoded the hippocampus development and evolution (Tosches, et al. 2018; Zhong, et al. 2020). It is worth noting that dozens of putative PSGs were related to cell types of both excitatory and inhibitory neurons, contributed to the connectivity and development of synapses and dendrites (Li, et al. 2014; Pendleton, et al. 2018).

In the present study, we collected single nuclei from the hippocampus of a Beagle dog and performed the comparative analysis of gene expression profiles of 105,057 nuclei. We defined 26 cell clusters and identified a set of marker genes for each cell type. Interestingly, 86 of differentially expressed genes (DEGs) were putative PSGs during dog domestication. The comprehensive cell landscape of the hippocampus could help us establish correspondence between cell types in nervous system and putative PSGs in dogs and facilitate the understanding of molecular features of cells during domestication.

## Results

### Single-nuclei transcriptomics of the dog hippocampus constructed using SPLiT-Seq

A standardized snRNA-seq pipeline was built using the SPLiT-seq (Rosenberg, et al. 2018). The scRNA-seq data of single nuclei from the hippocampus of a 5-month-old Beagle dog were obtained with the random and oligo(dT) primers. After detailed preprocessing and filtering (**Methods**), we created a digital expression matrix of 105,057 single nuclei with a median of 804 genes and 1,109 counts per nucleus. To remove the potential batch effects, partial principal component analysis (partial-PCA, **Methods**) was performed instead of the classical PCA and then uniform manifold approximation and projection (UMAP) was applied to project these two batches of transcripts into the common comparable two-dimensional (2D) space (**Figure S1A**). Furthermore, the Leiden community detection algorithm was employed to group these cells into 26 cell clusters (**Figure 1A**).

**Figure 1.**
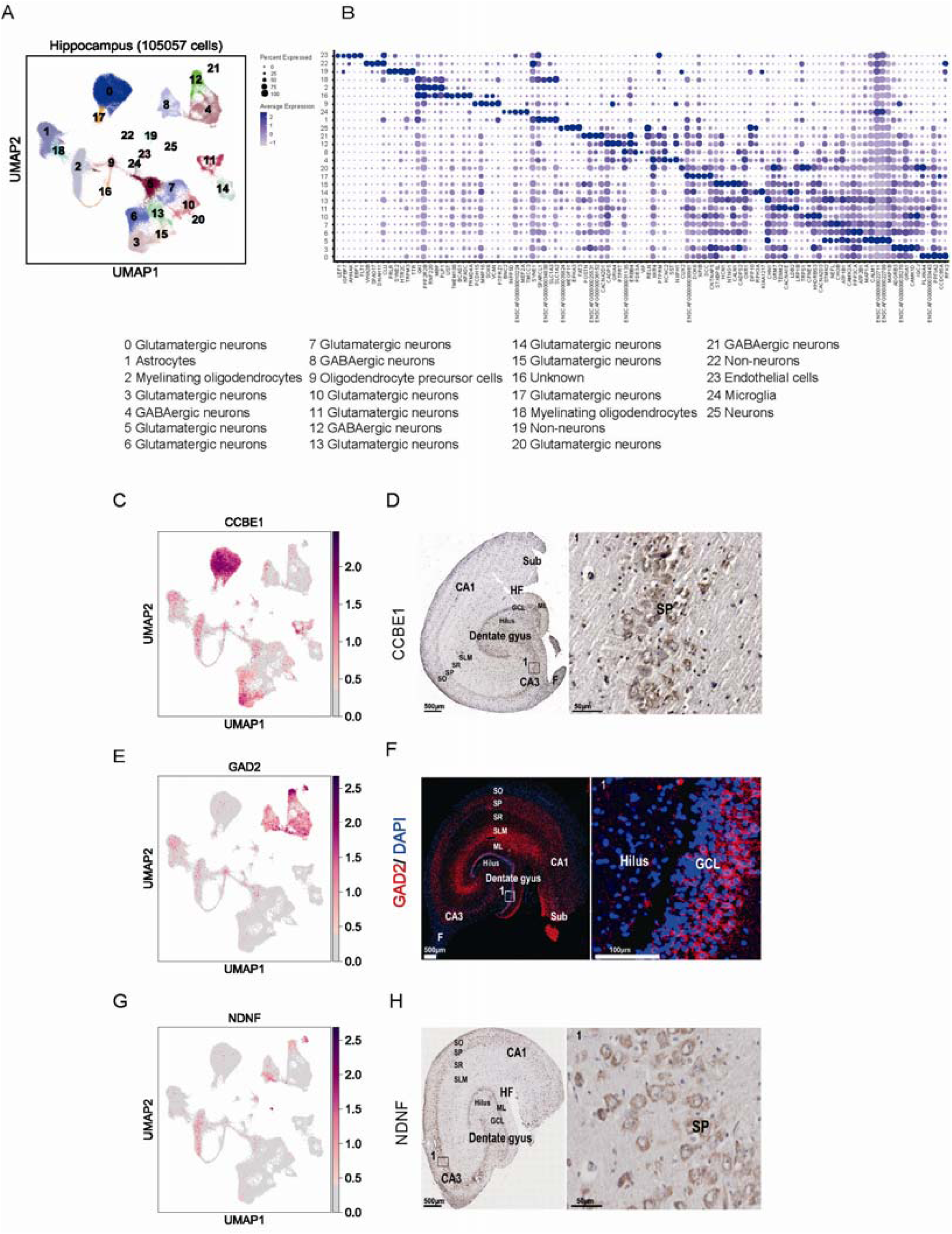
Single-nuclei transcriptomic landscape of dog hippocampus. (A) Clustering and visualization of transcriptomes of 105,057 single cells by UMAP. In this article, “cluster” means group of cells, is related with cluster analysis result. “Cell type” is group of clusters, we defined these clusters were same cell types. (B) Dot plot showing the averaged DEG expressions (z-cores) in different clusters (**Table S10**). (C) Visualization of *CCBE1* by UMAP. (D) *CCBE1* expressed in dog hippocampus CA3 SP region revealed by immunohistochemical staining. (E) Visualization of *GAD2* by UMAP. (F) *GAD2* expression in dog hippocampus revealed by immunohistochemical staining. Red: *GAD2*; Blue: DAPI; White square indicates higher magnification view of top region boxed. (G) Visualization of *NDNF* by UMAP. (H) *NDNF* expression in dog hippocampus revealed by immunohistochemical staining. Red point means cells were found differentially express DEGs. DG: Dentate Gyrus, GCL: Granule Cell Layer, SGZ: Subgranular Zone, ML: Molecular Layer, CA: Cornu Amonis, SO: Stratum Oriens, SP: Stratum Pyramidale, SR: Stratum Radiatum, SLM: stratum lacunosum-moleculare, Sub: Subiculum, HF: Hippocampus Fissure, A: Alveus.

The DEGs (p-value < 0.001 and log2fold-change > 0.25) of each cell cluster were used to assign most of cells to specific cell types, based on the known markers in mouse and human (Zeisel, et al. 2015; Rosenberg, et al. 2018; Zhang, et al. 2019) (**Table S1** and **Figure S1B**). Glutamatergic neurons, GABAergic neurons, Cajal-Retzius cell, oligodendrocyte precursor cells (OPCs), myelinating oligodendrocytes, endothelial cells, astrocytes, and microglial cells were detected in the Beagle dog hippocampus (**Figure 1B, Figure S1C, D**). They are also typical cells in the mammalian hippocampus.

Among the clusters, 12 ones (0, 3, 5, 6, 7, 10, 11, 13, 14, 15, 17 and 20) were identified as the glutamatergic neurons, which produce the most common excitatory neurotransmitter in the central nervous system (Baude, et al. 2009). Although cluster 3, 5 and 6 have a similar expression pattern, there were unique DEGs expressed in each cluster. For instance, both cluster 3 and 6 have high expression of *GRM7, TENM2*, and *CACNA1E*, while cluster 3 has no expression of *STMN2* and *NEFL* (**Figure 1B**). The *CCBE1* is a marker gene specifically in cluster 0 (**Figure 1C**). It marked the canine hippocampus CA3, SP (**Figure 1D**) the same as that in the mouse (© 2020 Allen Institute for Brain Science. Allen Brain map. Available from: http://mouse.brain-map.org/experiment/show/74513995). Eight glutamatergic neurons clusters (cluster 0, 5, 6, 7, 10, 13, 14, 20) were involved in the regulation of trans-synaptic signaling (GO:0099177, **Table S2**). Seven clusters (cluster 5, 6, 7, 13, 14, 17, 20) were involved in the plasma membrane bounded cell projection morphogenesis (GO:0120039, **Table S2**). The GO enrichment analysis showed that cluster 7 and 13 were enriched in cognition (GO:0050890, **Table S2**), cluster 11 and 20 were enriched in behavior (GO:0007610, **Table S2**), and cluster 13 were enriched in adult locomotory behavior (GO:0008344, **Table S2**).

Four clusters (cluster 4, 8, 12, and 21) are GABAergic neurons relating to a kind of inhibitory neurotransmitters in the central nervous system, indicated by the common marker gene *GAD2* (Zhang, et al. 2019) **(Figure 1E)**. Immunofluorescence (IF) analysis showed that the *GAD2* was highly expressed in the GCL and ML of DG area, and in SLM and SP of CA area (**Figure 1 F**). The GO enrichment analysis illustrated that four GABAergic neurons clusters enriched in the behavior (GO:0007610, **Table S2**) and the anterograde trans-synaptic signaling (GO:0098916, **Table S2**). Cluster 25 was identified as the Cajal-Retzius cell which is the project manager in the cerebral cortex and played a major role in the cortical development (Ogawa, et al. 1995; Villar-Cervino and Marin 2012; Meyer, et al. 2019). The marker gene of the cell type is *NDNF* (**Figure 1G, H**), which is also expressed in the Cajal-Retzius cells in human and mouse (Fan, et al. 2018; Rosenberg, et al. 2018). Beyond neuron cell clusters described above, another five clusters were representing non-neuron cells. Cluster 19 and 22 are two types of non-neuron cells. Cluster 1, 23, and 24 were identified as the astrocytes, endothelial cells, and microglial cells, respectively.

### Putative trajectory analysis reveals oligodendrocyte development in dogs

With the defined cell-type markers, cluster 2 and cluster 9 were inferred as myelinating oligodendrocytes and OPCs respectively (**Figure 2A**). Consistent with the classification, *CNP*, a myelin-related marker gene of cluster 2 (Yu, et al. 1994; Darmanis, et al. 2015), was high expressed in the Hilus, ML, SLM, F and A areas (**Figure 2B**). The *AGAP1* was a DEG in cluster 9 and could be used as a new marker of OPCs (**Figure 2B**). Cluster 16 was linked in cluster 2 and 9 **(Figure 1A**), with few specifically expressed genes to assign its identity. Therefore, we inferred that cluster 2, 9, 16 might be a development trajectory from OPCs to myelinating oligodendrocytes as the Alexander B. Rosenberg et al. 2018 found in the mouse hippocampus (Rosenberg, et al. 2018).

**Figure 2.**
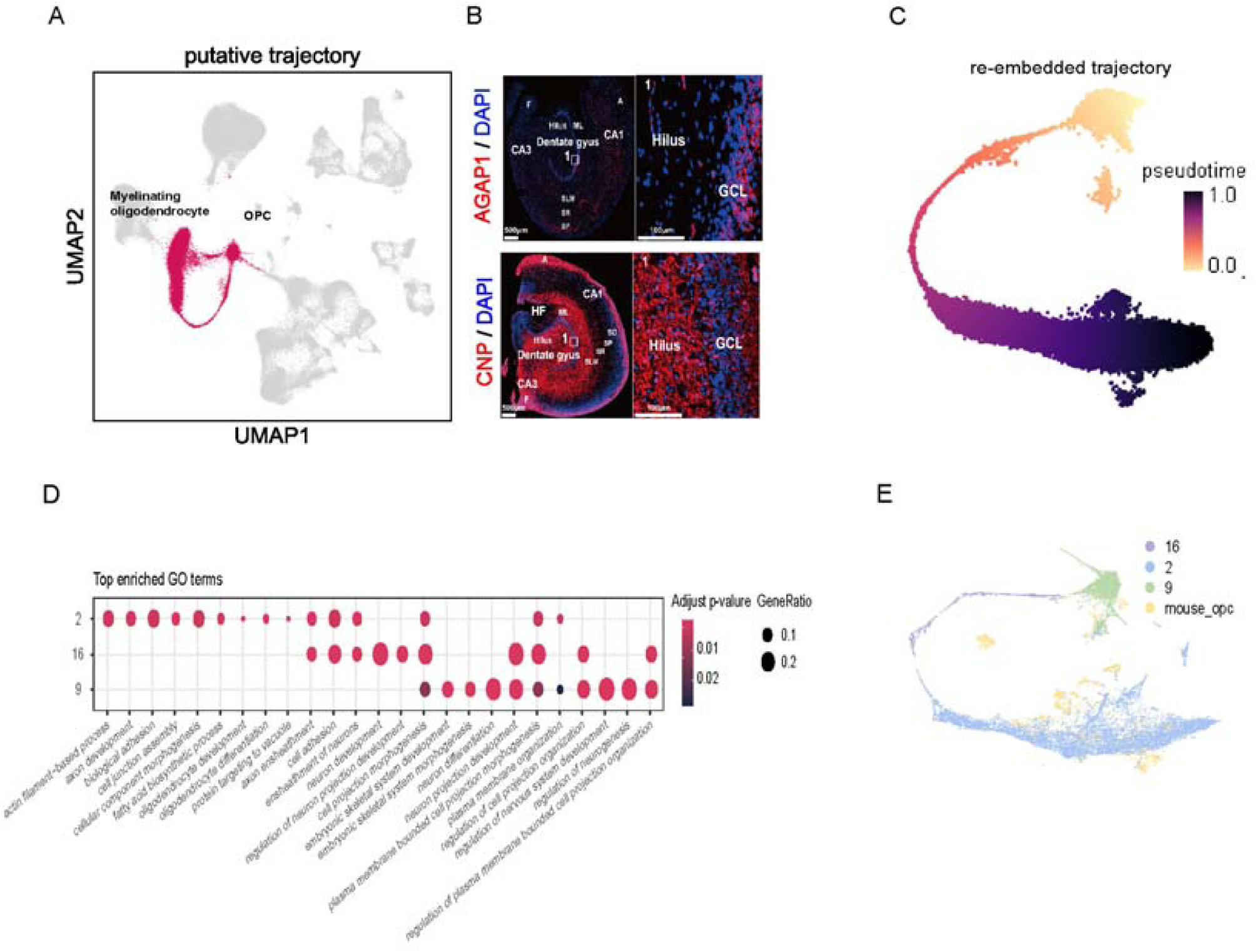
Putative trajectory analysis and cross-species transcriptome comparison between dog and mouse oligodendrocyte. (A) UMAP embedding of oligodendrocyte subclusters. Red indicates cells of cluster 2, 16, and 9. (B) Staining of AGAP1 and CNP of dog brain at hippocampus. Red: AGAP1 (top) and CNP (bottom), Blue: DAPI; White square indicates higher magnification view of top region boxed. (C) UMAP embedding of OPCs putative trajectory. Cells are colored by pseudotime, with dark colors representing mature cell stages and light colors representing immature cell stages. (D) GO enrichment analysis of OPCs differentiation clusters. Color bar means adjust p-value; dot size means gene ratio. (E) UMAP visualization of 20,989 single cells (15,920 from dog and 5,069 from mouse) analyzed by LIGER, color-coded by cell types.

To validate the hypothesis, we constructed the single-nuclei trajectories using the Slingshot (Street, et al. 2018) based on the re-calculated UMAP embedding and the original cluster labels (**Figure 2C**). As result, *PTPRZ1*, the marker gene for precursor cells that maintained OPCs in an undifferentiated state (Kuboyama, et al. 2012), was expressed in cluster 9; *SOX6*, was expressed in cluster 16 and repressed oligodendrocyte differentiation (Stolt, et al. 2006); *PLP1* is another myelin-related gene expressed in cluster 2 (**Figure S2A**). The GO enrichment analysis showed that oligodendrocytes (cluster 2, 9 and 16) were involved in different biological pathways (**Figure 2D**). The DEGs in cluster 9 (OPCs) were related to neuron differentiation (GO:0030182, **Table S2**) and regulation of neurogenesis (GO:0050767, **Table S2**). Those in cluster 16 were related to cell projection morphogenesis (GO:0048858, **Table S2**) and neuron projection development (GO:0031175, **Table S2**). Those in cluster 2 were enriched in ensheathment of neurons (GO:0007272, **Table S2**) and axon ensheathment (GO:0008366, **Table S2**). The DEGs of the putative trajectory were consistent with the differentiation process of oligodendrocyte development, and we also verified this result on mouse data (**Figure 2E, Figure S2B-E**).

### Significant convergence between DEGs and putative PSGs in domestication

Previous studies showed that dog putative PSGs were enriched in the neurological process such as learning and memory (Freedman, et al. 2016; Wang, et al. 2016) and expressed specifically in brain tissues (Li, et al. 2013). To identify which type of cell plays an important role during domestication, we gathered the DEGs in the 26 clusters (with p-value < 0.001 and log2fold-change > 0.25) and compared them with the putative PSGs from published studies (vonHoldt, et al. 2010; Axelsson, et al. 2013; Freedman, et al. 2016; Wang, et al. 2016; Pendleton, et al. 2018). The genes expressed in more than three cells on the dog hippocampus were used as the background (**Methods**). The results showed that 630 of 841 putative PSGs were detected in the hippocampus single nuclei transcriptomes (**Table S3, Figure S3**). 86 of 630 detected putative PSGs occurred in the 1,628 DEGs with a statistical significance of 5.10E-09 (**Table 1**). However, the 1,000 random gene sets with equal size are not significant with the p-value of 0.47. Since Freedman et al. 2016 did a statistical analysis to determine the likelihood that specific genes were under selection, we used their putative PSGs to verify the statistical significance. As result, 16 genes occurred in both DEGs and putative PSGs with the p-value of 6.99E-03. Furthermore, there is also a significant overlap between DEGs and the putative PSGs reported at least in two published studies (p-value of 2.45E-07, **Table S4**).

**Table 1.**
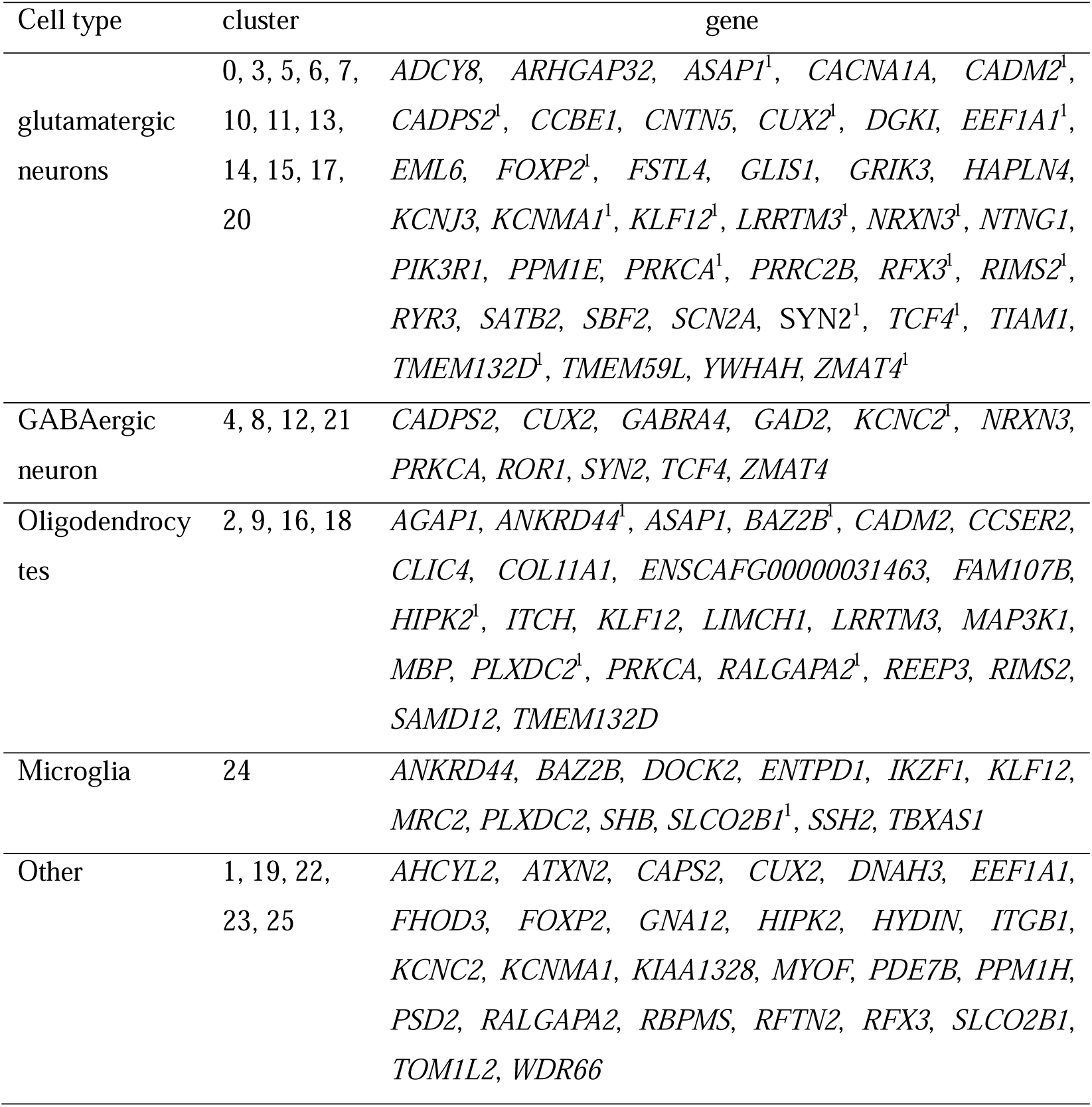
Overlapped genes between putative PSGs and DEGs. ^1^indicates this gene appears in different cell types, i.e., *CUX2* in Glutamatergic neurons, GABAergic neurons and non-neurons.

To explore whether putative PSGs were of higher or lower specificity across clusters or cell types, we calculated the entropy of putative PSGs. The higher entropy-specificity score for a gene implies its uniqueness in specific clusters or cell types (Martinez and Reyes-Valdes 2008). We found that the putative PSGs occurred in more than one literature have significantly higher entropy specificity across both clusters and cell types compared with the background (**Table 2**). To figure out which cluster is highly enriched with putative PSGs, we did enrichment analysis for each cluster-specific in the putative PSG set (**Table 3**). Four clusters (cluster 2, 9, 14, 24) were significantly enriched in three putative PSGs sets, six clusters (cluster 0, 8, 11, 16, 17, 20) were enriched in two putative PSGs sets (with p-value <0.05).

**Table 2.** The entropy of putative PSGs in different cell types and cluster. ^2^bg: background genes that were expressed by at least a proportion of “proportion-cut” in any cell cluster to filter out those genes having low expression frequencies. PSG-841: putative PSGs in five articles; PSG-78: putative PSGs occurred in no less than two articles; PS-145: putative PSGs occurred in Freedman et al. 2016.

|  |  | Gene-specificity based on the average<br>expressions of cluster |  |  | Gene-specificity based on the average<br>expressions in a cell type |  |  |
| --- | --- | --- | --- | --- | --- | --- | --- |
|  | proportion-cut | PSG-841 | PSG-78 | PSG-145 | PSG-841 | PSG-78 | PSG-145 |
| PSG | 0 | 1.57E-01 | 1.46E-01 | 1.85E-01 | 2.11E-01 | 1.94E-01 | 2.36E-01 |
| bg | 0 | 2.47E-01 | 2.45E-01 | 2.45E-01 | 3.31E-01 | 3.28E-01 | 3.28E-01 |
| PSG | 0.1 | 7.96E-02 | 1.26E-01 | 1.05E-01 | 1.01E-01 | 1.52E-01 | 1.27E-01 |
| bg | 0.1 | 9.36E-02 | 9.28E-02 | 9.29E-02 | 1.20E-01 | 1.19E-01 | 1.19E-01 |
| PSG | 0.2 | 7.43E-02 | 1.23E-01 | 8.97E-02 | 9.21E-02 | 1.52E-01 | 1.10E-01 |
| bg | 0.2 | 8.97E-02 | 8.87E-02 | 8.89E-02 | 1.14E-01 | 1.13E-01 | 1.13E-01 |
| PSG | 0.25 | 7.00E-02 | 1.07E-01 | 9.32E-02 | 8.68E-02 | 1.34E-01 | 1.15E-01 |
| bg | 0.25 | 9.00E-02 | 8.88E-02 | 8.89E-02 | 1.14E-01 | 1.13E-01 | 1.13E-01 |

**Table 3.** Enrichment analysis for each cluster-specific genes in putative PSG sets. PSG-841: putative PSGs in five articles; PSG-78: putative PSGs occurred in no less than 2 articles, PS-145: putative PSGs occurred in Freedman et al. 2016.

| cluster | PSG-841 | PSG-78 | PSG-145 |
| --- | --- | --- | --- |
| 0 Glutamatergic neurons | 4.06E-05 | 6.18E-03 | 1.42E-01 |
| 0 Glutamatergic neurons(random) | 4.33E-01 | 2.32E-01 | 3.00E-01 |
| 3 Glutamatergic neurons | 2.38E-03 | 5.35E-02 | 4.86E-01 |
| 3 Glutamatergic neurons(random) | 4.30E-01 | 2.25E-01 | 3.04E-01 |
| 5 Glutamatergic neurons | 4.37E-01 | 3.76E-01 | 2.04E-01 |
| 5 Glutamatergic neurons(random) | 4.27E-01 | 2.63E-01 | 3.26E-01 |
| 6 Glutamatergic neurons | 9.51E-02 | 2.31E-01 | 3.75E-01 |
| 6 Glutamatergic neurons(random) | 4.01E-01 | 1.87E-01 | 2.63E-01 |
| 7 Glutamatergic neurons | 1.38E-01 | 2.57E-01 | 4.11E-01 |
| 7 Glutamatergic neurons(random) | 4.06E-01 | 1.94E-01 | 2.66E-01 |
| 10 Glutamatergic neurons | 1.02E-02 | 2.77E-01 | 4.39E-01 |
| 10 Glutamatergic neurons(random) | 4.10E-01 | 2.14E-01 | 2.89E-01 |
| 11 Glutamatergic neurons | 7.28E-05 | 2.43E-02 | 6.88E-02 |
| 11 Glutamatergic neurons(random) | 4.22E-01 | 1.74E-01 | 2.43E-01 |
| 13 Glutamatergic neurons | 1.83E-01 | 2.15E-01 | 3.50E-01 |
| 13 Glutamatergic neurons(random) | 4.08E-01 | 1.74E-01 | 2.52E-01 |
| 14 Glutamatergic neurons | 1.77E-02 | 2.04E-04 | 1.51E-02 |
| 14 Glutamatergic neurons(random) | 4.04E-01 | 1.99E-01 | 2.69E-01 |
| 15 Glutamatergic neurons | 5.16E-02 | 2.52E-01 | 4.05E-01 |
| 15 Glutamatergic neurons(random) | 4.11E-01 | 1.95E-01 | 2.79E-01 |
| 17 Glutamatergic neurons | 1.54E-03 | 3.47E-02 | 4.08E-01 |
| 17 Glutamatergic neurons(random) | 4.05E-01 | 1.98E-01 | 2.81E-01 |
| 20 Glutamatergic neurons | 8.16E-02 | 4.23E-02 | 2.06E-02 |
| 20 Glutamatergic neurons(random) | 4.33E-01 | 2.12E-01 | 2.79E-01 |
| 4 GABAergic neurons | 7.88E-02 | 4.18E-03 | 1.13E-01 |
| 4 GABAergic neurons(random) | 4.05E-01 | 2.14E-01 | 2.91E-01 |
| 8 GABAergic neurons | 2.32E-03 | 2.09E-03 | 3.64E-01 |
| 8 GABAergic neurons(random) | 3.99E-01 | 1.80E-01 | 2.53E-01 |
| 12 GABAergic neurons | 2.08E-01 | 4.87E-03 | 4.58E-01 |
| 12 GABAergic neurons(random) | 4.18E-01 | 2.21E-01 | 2.99E-01 |
| 21 GABAergic neurons | 1.57E-01 | 5.90E-02 | 5.06E-01 |
| 21 GABAergic neurons(random) | 4.27E-01 | 2.39E-01 | 3.05E-01 |
| 25 Cajal-Retzius | 2.69E-01 | 1.78E-01 | 2.95E-01 |
| 25 Cajal-Retzius(random) | 3.79E-01 | 1.48E-01 | 2.19E-01 |
| 1 Astrocytes | 1.46E-01 | 1.07E-01 | 2.62E-01 |
| 1 Astrocytes(random) | 4.26E-01 | 2.81E-01 | 3.50E-01 |
| 24 Microglia | 1.09E-05 | 3.34E-04 | 3.11E-04 |
| 24 Microglia(random) | 4.15E-01 | 2.13E-01 | 2.82E-01 |
| 9 Oligodendrocyte precursor cells | 2.38E-03 | 5.30E-04 | 5.45E-04 |
| 9 Oligodendrocyte precursor cells(random) | 4.42E-01 | 2.21E-01 | 3.07E-01 |
| 16 Unknown | 8.92E-02 | 2.72E-03 | 3.84E-03 |
| 16 Unknown(random) | 4.42E-01 | 2.84E-01 | 3.55E-01 |
| 2 Myelinating oligodendrocytes | 6.22E-03 | 2.57E-02 | 5.62E-03 |
| 2 Myelinating oligodendrocytes(random) | 4.42E-01 | 2.96E-01 | 3.53E-01 |
| 18 Myelinating oligodendrocytes | 3.81E-01 | 4.52E-02 | 1.22E-01 |
| 18 Myelinating oligodendrocytes(random) | 4.05E-01 | 2.18E-01 | 2.88E-01 |
| 19 Non-neuron | 1.46E-01 | 2.37E-03 | 2.62E-01 |
| 19 Non-neuron(random) | 4.45E-01 | 2.87E-01 | 3.46E-01 |
| 22 Non-neuron | 2.12E-01 | 1.59E-01 | 7.21E-01 |
| 22 Non-neuron(random) | 4.43E-01 | 3.11E-01 | 3.71E-01 |
| 23 Endothelial cells | 2.28E-01 | 1.01E-01 | 7.12E-02 |
| 23 Endothelial cells(random) | 4.26E-01 | 2.77E-01 | 3.45E-01 |

Glutamatergic receptors constitute major excitatory transmitter system, and dog excitatory synaptic plasticity increased in the domestication process (Li, et al. 2014). 40 of 86 putative PSGs were expressed in the glutamatergic neurons, 28 in cluster 0, 11, 14, 17 and 20. The GO enrichment analysis showed that DEGs of cluster 0 enriched in the regulation of cell migration (GO:0030334, **Table S2**) and regulation of cell motility (GO:2000145, **Table S2**), those in cluster 11 and cluster 14 involved in neuron differentiation (GO:0030182, **Table S2**), and those in cluster 17 and cluster 20 enriched in axon guidance (GO:0007411, **Table S2**) and neuron projection guidance (GO:0097485, **Table S2**). *CUX2* was highly expressed in DG granular cells as glutamatergic neurons DEG in cluster 11 and 20, and it was also expressed in CA granular cells, the same expression pattern in the mouse (**Figure 3** and **Figure S4B**) (Yamada, et al. 2015). Another putative PSG, *GRIK3* in cluster 14 was detected mainly in GCL of DG and SP of CA area (**Figure 3**). It is worth noting that putative PSGs were significantly enriched in cluster 14 (**Table 3**), and the DEGs in cluster 14 were enriched in synapse organization (GO:0050808, **Table S2**), neuron differentiation (GO:0030182, **Table S2**), synaptic signaling (GO:0099536, **Table S2**), and synapse assembly (GO:0007416, **Table S2**), It suggests that these clusters may involve in stress responses (O’Rourke and Boeckx 2020).

**Figure 3.**
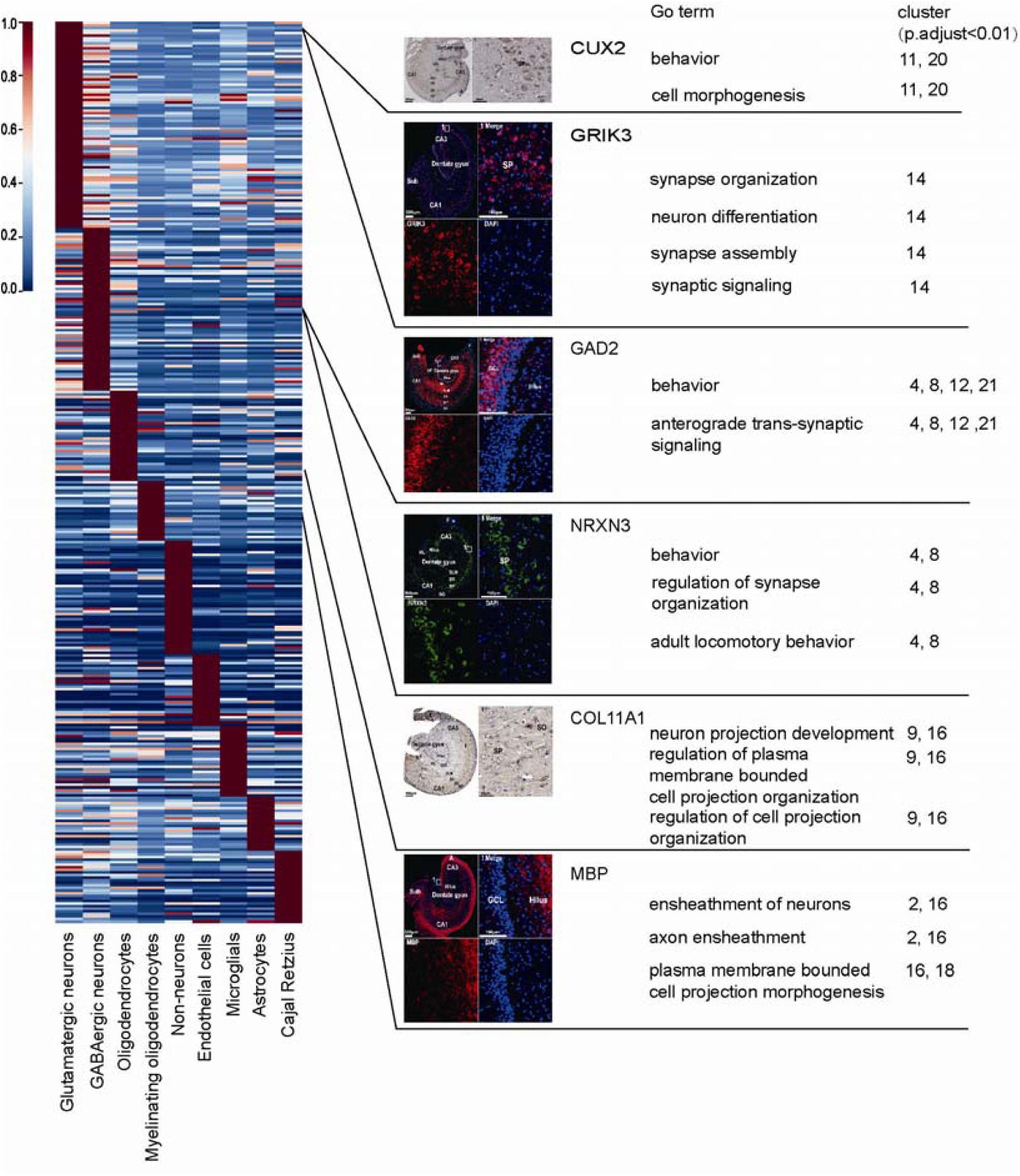
The mean expression of DEGs in different cell types. The values of each gene (row) were its original ones divided by its maximum. Six genes on the right of the picture were differentially expressed in different cell types and the GO enrichment analysis showed their relevant gene functions. It is worth to note that *CUX2* is a DEG in glutamatergic neuron as compared to other cells although highly expressed both in glutamatergic neuron and non-neurons. High-definition immunohistochemical staining is in the **Figure S4A**.

11 putative PSGs were expressed in the GABAergic neuron, and seven of them belong to cluster 8. GO enrichment analysis suggested DEGs in cluster 8 were involved in behavior (GO:0007610, **Table S2**), forebrain development (GO:0030900, **Table S2**) and adult locomotory behavior (GO:0008344, **Table S2**). *GAD2* as a candidate gene (**Figure 3**), to regulate brain development and behavior during domestication (Pendleton, et al. 2018), and it was expressed in cluster 4, 8, 12 and 21 (**Figure 1E**). Moreover, DEGs in cluster 4, 8, 12 and 21 enriched with behavior (GO:0007610, **Table S2**) and anterograde trans-synaptic signaling (GO:0098916, **Table S2**). Previous studies suggested that ablation GAD2, mouse reduced freezing and increased flight and escape behavior (Stork, et al. 2003). *NRXN3* (cluster 4 and 8) encodes a receptor and cell adhesion molecule in the nervous system, and it was highly expressed in the SP of the hippocampus CA area (**Figure 3**). GO enrichment analysis showed DEGs in cluster 4 and 8 were relevant to behavior (GO:0007610, **Table S2**), regulation of synapse organization (GO:0050807, **Table S2**), and adult locomotory behavior (GO:0008344, **Table S2**). A previous study also confirmed that putative PSGs in the Chinese native dogs enriched in locomotory behavior (Li, et al. 2013).

24 putative PSGs occurred in the oligodendrocytes including cluster 2, 9, 16, 18. Three of them are related to OPC differentiation. Oligodendrocytes are a kind of cells that primarily form the central nervous system (CNS) myelin. Myelination is critical for the normal functioning of the CNS (Fancy and Miller 2020). One of the overlapped genes *COL11A1* (cluster 9, 16) marked OPCs and was expressed in the partial area of hippocampus CA (**Figure 3**). Previous studies showed that the OPC was important for human white matter expansion and myelination (Huang, et al. 2020). Electron microscopic analysis revealed that glutamatergic synapses connected OPCs and neurons (Bergles, et al. 2000). Myelinating oligodendrocyte was the result of OPC differentiation and made saltatory conduction of nerve impulses possible (Rivers, et al. 2008; Freeman and Rowitch 2013). *MBP* was a myelinating oligodendrocyte cell marker, expressed in cluster 2, 16 and 18, and highly expressed in the Hilus, ML, SLM and other hippocampus regions (**Figure 3**). As a domestication gene, *MBP* encoded myelin basic protein. Domestication influenced myelination and/or axonal diameter in rabbits (Brusini, et al. 2018), and affected neurological function in the dogs (Freedman, et al. 2016). The relevant functions on these clusters were ensheathment of neurons (cluster 2, 16, GO:0007272, **Table S2**), plasma membrane bounded cell projection morphogenesis (cluster 16, 18, GO:0120039, **Table S2**), and axon ensheathment (cluster 2, 16, GO:0008366, **Table S2**).

### Reconstructed gene regulatory networks of dog hippocampus

We reconstructed dog hippocampus gene regulatory networks by GENIE3 software (Huynh-Thu, et al. 2010). To attenuate the effects of noise and outliers, we used 4,523 genes and 4,281 pseudo cells (**Methods** about WGCNA), which contained all the cell types in this study.More than two million interactions were found between the putative regulatory genes and target genes with 2,239 putative regulatory genes were in the top 1% (**Table S5**). The transcription factors in dogs (Canis familiaris) from AnimalTFDB3.0 (Hu, et al. 2019) were used to verify these putative regulatory genes. As the results shown, there were 118 putative regulatory genes, and 11 of those are both PSGs and DEGs, including Cut Like Homeobox 2 (*CUX2*) (Freedman, et al. 2016; Pendleton, et al. 2018) and Regulatory Factor X3 (*RFX3*) (Wang, et al. 2016) in the glutamatergic neurons. GO enrichment analysis showed that target genes of *CUX2* were related to glutamatergic synaptic transmission (GO:0035249, **Table S6**). The transcription factor CUX2 is involved in early neocortical circuits, cellular fate selection and mechanosensation development (Bachy, et al. 2011; Miskic, et al. 2021). Another regulatory gene *RFX3* was enriched in neurogenesis (GO:0022008, **Table S6**), neuron development (GO:0048666, **Table S6**) and locomotion (GO:0040011, **Table S6**). Taken together, results above implied that glutamatergic neurons may involve in dog behaviors adaptive evolution.

## Discussion

In this study, we presented the single-nuclei transcriptomics of the dog hippocampus and identified 26 cell clusters based on 105,057 single-nuclei transcriptomes, such as glutamatergic neurons, GABAergic neurons, Cajal-Retzius cells, oligodendrocyte precursor cells (OPCs), myelinating oligodendrocytes, endothelial cells, astrocytes, and microglial cells. Besides, our study demonstrated the trajectory of oligodendrocytes differentiation based on the re-calculated UMAP embedding analysis.

As a subtype of inhibitory neurons, the GABAergic neuron has been linked to the response to learning and fear memory (Harris and Westbrook 1998; Stork, et al. 2002). Reduced fear and aggression are important traits selected by humans in the first step of animal domestication, which help animals not only live commensally with humans, but also stay in a crowded environment (de Kloet, et al. 2005; Price 2008). Especially, *GAD2* could influence fear behavior and relieve sensitized pain behavior (Stork, et al. 2003; Zhang, et al. 2011), *KCNC2*(Kv3.2) expressed in the GABAergic neurons synaptic which had been related to diurnal and circadian rhythms of wheel-running behavior (Chow, et al. 1999; Kudo, et al. 2011).

Glutamate receptors play an important role in the central nervous system, responded for basal excitatory synaptic transmission, some of them are involved in learning and memory, and there were great changes of glutamate receptors in animal domestication and modern human evolution (O’Rourke and Boeckx 2020). *FOXP2* is a DEG in cluster 14. A previous study illustrated that laboratory rats are good at learning than wild rats because of the selected *FOXP2* (Zeng, et al. 2017). The FOXP2 protein in humans and mice is extremely conserved (Enard, et al. 2002). Putative target genes of the FOXP2 showed these genes play roles in the CNS patterning, development, and function (Fisher and Scharff 2009). The absence of FOXP2 protein could lead to developmental delays, severe motor dysfunction, and neural abnormalities in mice (French, et al. 2007). Besides *FOXP2*, another two putative PSGs in glutamatergic neurons also need to note. One is *GRIK3*. Genome wide studies showed that signals of selection exist in the dog (Axelsson, et al. 2013; Freedman, et al. 2016), cattle (Qanbari, et al. 2014) and sheep (Naval-Sanchez, et al. 2018). *GRIK3* may play important roles in reducing stress responses between domesticated animals and humans, leading to more prone to human custody (O’Rourke and Boeckx 2020). Another is *GRM8* (mGluR8) as one of the DEGs in cluster 14 and one of the parallel evolutionary genes between humans and dogs (Wang, et al. 2013). As a member of the Group III receptor subunit, mice lacking GRM8 are heavier, reduce locomotory behavior, increase anxiety-like behaviors in the open field (Duvoisin, et al. 2005; Duvoisin, et al. 2010).

Furthermore, DEGs in cluster 14 were found enriched in regulation of synapse structure or activity (GO:0050803, **Table S2**) and regulation of nervous system development (GO:0051960, **Table S2**). It revealed that cells in cluster 14 may play an important role in a decreased stress response not only in domesticated animals, but also in humans (O’Rourke and Boeckx 2020). These references could verify the hypothesis that glutamatergic neurons changed the behavior during domestication and reduce fear response for dogs through regulating cells in the hippocampus. It suggested that cluster 14 is probably an important cluster in dog domestication.

Genome scan for selection signature in domestic animals also revealed that putative PSGs enriched in the nervous system development (Montague, et al. 2014; Qanbari, et al. 2014; Schubert, et al. 2014; Qiu, et al. 2015). Here we found the putative PSGs in domestication were involved in the development of oligodendrocytes. For instance, *MBP*, a putative PSG that may lead to behavioral differences between dogs and wolves (Cagan and Blass 2016), was also one of the marker genes of myelinating oligodendrocytes. The function is to encode the myelin basic protein, a component of the myelin sheath, considered to be essential for saltatory impulse propagation (chiron and Miron 2007; Simons and Nave 2016). In addition, domestication may reduce myelination level, compromise neural conduction, and change the size of brain structure relevant for memory, reflexes, and fear processing (Brusini, et al. 2018). The lower reactive level is a component of domestication syndrome that is a collection of common traits in domestic animals (Wilkins, et al. 2014).

It is worth to note that the PSGs from the published studies were described as putative PSGs in the present study. Wang et al., vonHoldt et al., Axelsson et al., and Pendleton et al. reported 205, 29, 122 and 429 PSGs through the whole genome scan for selection, respectively (vonHoldt, et al. 2010; Axelsson, et al. 2013; Wang, et al. 2016; Pendleton, et al. 2018). None of those were validated as the PSGs by the additional statistical analysis nor the experimental assays. Freedman et al. reported 145 PSGs and performed the statistical analysis to determine the likelihood under selection (Freedman, et al. 2016). Thus, we performed statistical analysis using the putative PSGs in Freedman et al 2016 only, and those reported at least in the two references, leading to the same significant overlap. Interestingly, *GRIK3* in cluster 14 was reported in four published studies, except vonHoldt, et al. 2010. Glutamate plays a major role in the excitatory transmitter system, responded for basal excitatory synaptic transmission which is involved in learning and memory, hypothalamic-pituitary-adrenal activity, and tameness and the reduction of reactive aggression (Herman, et al. 2004; Baude, et al. 2009;O’Rourke and Boeckx 2020). Nevertheless, further statistics or experiment assays should be done in the future to verify the status of the PSGs.

In conclusion, 105,057 single-nuclei transcriptomes were classified into 26 clusters in this study and were defined with distinct identities. Further exploration revealed 86 genes that were overlapped between putative PSGs and DEGs. Besides, we illustrated the OPC differentiation trajectory based on the UMAP embedding. Our results contributed to defining the cell types and revealing the development of oligodendrocyte in the dog hippocampus, illustrating the difference between the sub-cell types and connecting the gap of our understanding between the molecular and cellular mechanisms of animal domestication.

## Materials and Methods

### Ethics approval and consent to participate

Hippocampus tissue was collected from a five-month-old Beagle ordered from the department of laboratory animal science, Kunming Medical University. All the animal processing procedures and experiments performed in the present study were approved by the Animal Ethics Committee of Kunming Institute of Zoology, Chinese Academy of Sciences (SMKX-20160301-01).

### Tissue dissection and nucleus extraction

Broader dissections (no layer enrichment or multiple layers combined) were used to facilitate isolation of sufficient number of cells. Nucleus extraction protocol was adapted from Rosenberg et al. 2018 (Rosenberg, et al. 2018). Briefly, a dounce homogenizer (Wheaton, cat. no. 357538) was used for nucleus extraction. Hippocampus was homogenized in the dounce homogenizer (Kimble) 10X with lose pestle and 20X with tight pestle. A homogenization buffer (4.845 mL of NIM1 buffer (250 mM sucrose, 25 mM KCl, 5 mM MgCl2, 10 mM Tris pH=8.0), 5 _μ_L of 1 mM DTT, 50 _μ_L of Enzymatics RNase Inibitor (40U/_μ_L), 50 _μ_L of SuperaseIn RNase Inhibtor (20U/_μ_L), 50 _μ_L of 10% Triton-X100) was used to make a homogeneous nucleus solution and protect the RNA from degradation. Nuclei were collected by the centrifugation and filtered through sterile 40_μ_m cell strainer (Corning) into 1.5ml tubes. The nucleus concentration was checked by the hemocytometer to ensure it was within 1,000,000 nuclei/ml.

### Library preparation

SPLiT-seq (split-pool ligation-based transcriptome sequencing) was used to generate the libraries with Uniquely Barcoded Cells (UBC) (Rosenberg, et al. 2018). Briefly, nuclei were distributed into 48 individual wells in two 96-well plates. Each well loaded about 5,000-8,000 nuclei. The random hexamer and the anchored poly(dT)15 barcoded RT primers were respectively distributed into the 48 individual wells with nuclei of the 96-well. The links between well ID and barcode were recorded for downstream data processing. For the other protocol of the reverse transcription, ligation barcoding, lysis, cDNA amplification, tagmentation and Illumina amplicon generation were processed according to the SPLiT-seq protocol Version 3.0. The libraries were processed on the Illumina platform for sequencing of 150 bp pair-end reads.

### Processing of raw scRNA-seq data

According to the procedure of SPLiT-seq, the site of read 11-18 bp that did not match the barcode list were removed. The sequencing data was processed with the Drop-seq core computational protocol (“http://mccarrolllab.org/dropseq/”) to align, filter, and count unique molecular identifiers (UMIs) per sample by default. Data were mapped to the dog reference genome CanFam3.1 (GCA_000002285.2) and the transcriptome annotation from the Ensembl database, Canis_familiaris. CanFam3.1.92. Cells with more than 350 and no more than 5,000 expressed genes were retained. After that, genes expressed in less than three cells were filtered out. At the same time, cells that were expressed in a high proportion of mitochondrial genes (> 0.03) were also removed. As a result, 18,039 cells from the random group and 87,018 cells from the poly-T group were used for further analysis.

### Data normalization and HVG selectionmo

The gene-by-cell matrices were adjusted by a total-count normalization. Formally, denote *x*_*ij*_ as the raw count of gene *i* in cell *j*. For each cell *j*, counts were divided by its total count *c*_*j*_ = ∑_*i*_ *x*_*ij*_ of this cell and then multiplied by the median of all the total counts *m= median*{*c*_*j*_}. The total-size normalized expressions 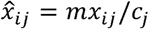 were then log-transformed to 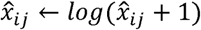 for downstream analysis. Highly variable genes (HVGs) were selected separately within random-group and poly-T-group, using the build-in function “scanpy.pp.highly_variable_genes” with the Python package ScanPy (Wolf, et al. 2018). Genes with a (log-normalized) mean expression above 0.025 and a normalized dispersion higher than 0.25 were identified as highly variable ones in each group. Finally, we took those genes that were highly variable in both groups and those with top dispersion as final HVGs for downstream analysis. This process resulted in 2,000 HVGs for downstream analysis.

### Dimensionality reduction and clustering

Before performing dimensionality reduction, the gene-by-cell expression matrix was centralized and scaled within each group of the same RNA capturing primer, for elimination of the batch effect. The partial-PCA was performed to combine the random and poly-T groups. That was calculating the principal components on the polyT-group first, and projecting the random-group onto the same PC space of the polyT-group. We selected the top 50 principal components (PCs) with the highest explained variances. We also tried different numbers of PCs and got similar results. UMAP (Mcinnes and Healy 2018) was used to embed each cell from the reduced PC space into a 2D space. It first computed the approximate k nearest neighbors (KNNs) for each data point, built a weighted mutual-KNN graph with each node representing each cell, and embedded each node of the graph into the low-dimensional space. We computed 30 approximate nearest neighbors for each single cell based on cosine distance in the PC space. Leiden community detection algorithm (Traag, et al. 2019) was applied (with resolution=0.8, which resulted a fine grained clustering) onto the weighted KNN graph built by UMAP to cluster single cells into distinct groups.

### Identification of DEGs and enrichment analysis

We compared the transcriptomic profile of each cluster versus the others using three differential analysis methods to get reliable DEG sets (Student’s t-test, Wilcox (Wilcoxin 1947), MAST (Finak, et al. 2015)) with the same thresholds (p-values less than 0.001 and log2fold-changes more than 0.25). The intersection of them were 64 to 233 DEGs for each cluster (**Table S7**). Each detected cluster was mapped to cell types or intermediate states by matching their corresponding DEGs to Allen Brain Atlas database and consulting published literatures. GO enrichment analysis of these DEGs was performed by the R package “clusterProfiler” (Yu, et al. 2012) with the reference database “org.Cf.eg.db” (Carlson 2019). Instead of the genes in whole genome of the dog, we used the genes that were detected in the present hippocampus data as the background.

### Categorization of genes using WGCNA

We adopted WGCNA (Weighted gene correlation network analysis) (Langfelder and Horvath 2008) analysis to detect gene modules with the R package “WGCNA” (R version 3.6.3, https://cran.r-project.org/web/packages/WGCNA; package version 1.69), which was initially designed for the bulk RNA-seq data. For the convenience of calculation, we used 4,523 genes expressed in more than 25% of populations in at least one cluster. Considering the large number of single-nuclei and the sparsity of the expression profiles, samples used in the WGCNA were created by aggregating and averaging each of the small clusters (< 100 cells) of single cells of similar transcripts, resulting 4,281 pseudo cells. The soft power value (power = 4) was determined by inspecting the soft-threshold-mean-connectivity curve plot. Modules with distance less than 0.25 (i.e., correlation more than 0.75) were merged. The module-cluster relationships were evaluated with the Pearson correlation coefficients between the module-memberships (MMs, a gene-by-module matrix) for genes and the gene-significances (GSs, a gene-by-cluster matrix) for clusters. The MM was defined as the Pearson correlation coefficients between the pseudo cell expressions and the eigengenes of the modules (computed by the WGCNA), while the GS was defined as the Pearson correlation coefficients between the pseudo cell expressions and the one-hot coded cluster labels.

### Putative trajectory analysis for a subset of hippocampus cells

To validate our hypothesis of the trajectory, we separated these clusters of cells, and redid dimensionality reduction using PCA and UMAP. Moreover, we analyzed the polyT group only for that they hold the majority (15,791 cells, nearly 90% of the entire trajectory) of the transcriptomes that formed the trajectory. Therefore, we can keep the biological information from being lost without introducing additional technical noise. In detail, we selected the top 20 PCs according to the PC-elbow plot, found 10 nearest neighbors for each cell using Euclidean distance and performed UMAP with the parameter “min_dist = 0.2”. After that, the pseudo-time of each cell on the trajectory was inferred using Slingshot, based on the re-calculated UMAP embeddings and the original cluster labels. The student’s t-test was performed to find out the DEGs within the three clusters that formed the trajectory.

### Cross-species integration of oligodendrocyte

Integration of single-cell data from mouse and domestic dogs was processed with LIGER (liger) (Welch, et al. 2019), which is an R package for integrating and analyzing multiple single-cell datasets. We downloaded 5,072 transcriptomes of single cells from the mouse oligodendrocyte lineage obtained at GEO (GSE75330) and separated cells of correlative clusters to do integrated analysis. We used LIGER with the following processing and parameters. First, we normalized the data to account for differences in sequencing depth and capture efficiency across cells, selected variable genes, and scaled the data with “var.thresh = 0.1”. Next, we performed integrative non-negative matrix factorization to identify shared and distinct metagenes across the datasets using the LIGER function “optimizeALS (k = 25)”. We performed a quantile alignment step “quantileAlignSNF” with the default settings and used UMAP to visualize the integrated data.

### Enrichment analysis of DEGs on putative PSG sets

The domesticated gene list was obtained from five studies (**Table S8**) (vonHoldt, et al. 2010; Axelsson, et al. 2013; Freedman, et al. 2016; Wang, et al. 2016; Pendleton, et al. 2018). We pooled the DEGs from all clusters for the enrichment analysis based on the hypergeometric test. The hypergeometric distribution, which describes the probability of k successes (random draws for which the object drawn has a specified feature) in n draws, without replacement, from a finite population of size N that contains exactly K objects with that feature, whereeach draw is either a success or a failure. The probability distribution function is given by 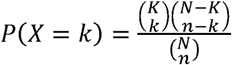, where 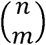 is the binomial coefficient. And the probability of over-representation is ∑_*k* ≤*i* ≤*n*_ *P*(*X*= *i*). In our case, we take a total of N=22,639 genes expressed in more than three cells as the gene universe instead of using the whole genome of the dog. K is the number of intersects between a given putative PSG set and the gene universe. We used the method to calculate not only all putative PSGs but also those only reported in Freedman et al. (2016) and in at least 2 references.

### Entropy specificities of putative PSG sets across clusters and cell-types

We followed (Martinez and Reyes-Valdes 2008) and computed the entropy specificity for each gene, based on the normalized average expressions, grouped by those 26 clusters and 10 major cell types. The computation was done by the build-in function ‘entropySpecificity’ of BioQC (Zhang, et al. 2017). To avoid the noises from the lowly expressed genes, we took the genes expressed in more than 10% of populations in at least one cluster as the background. The entropy-specificity scores were normalized and averaged for each putative PSG set and the backgrounds.

### Genetic regulatory networks (GRN) analysis by the GENIE3

Inferable regulatory links for each gene were predicted by the GENIE3 software (version 1.16.0, Huynh-Thu, et al. 2010) with the random forest machine learning algorithm. The links ranked by weight and only top 1% regulatory genes were reserved for subsequent analysis. Transcription factors of the dog were downloaded from the AnimalTFDB 3.0 (Hu, et al. 2019). GO enrichment analysis for target genes was performed using the g:Profiler (version e104_eg51_p15_3922dba) by the dog annotation (Raudvere, et al. 2019).

### Immunofluorescent (IF) and immunohistochemistry (IHC) staining

Formalin-fixed and paraffin-embedded hippocampus tissue specimens were sliced into 9 µm sections and then deparaffinized in turpentine oil (TO), rehydrated through graded ethanol solutions and antigen retrieval was performed by placing the slides at 95°C for 40 min in a microwave oven and allowed to cool at room temperature. The slides were washed three times with PBS, and then the slides were permeabilized with 1% Triton X solution for 10 min. After nonspecific binding was blocked by 5% Bovine serum albumin (BSA) for 1h at room temperature, the slides were incubated with primary antibodies overnight at 4 °C. For immunofluorescent staining, DAPI (4-6-diamidino-2-phenylindole) and DyLight 488-or DyLight 555-labeled secondary antibodies (1:400, Thermo Fisher Scientific, Waltham, MA) were added for 1.5h at room temperature. For immunohistochemistry staining, the slides were treated with HRP-labeled second antibody (Thermo Fisher Scientific) for 30 min. The slides were incubated with DAB until desired stain intensity is observed and then followed by slight hematoxylin counterstaining. The slides were finally dehydrated and mounted with a cover glass. Stained slides were visualized and imaged using a laser scanning confocal microscope (Olympus, Tokyo, Japan). Primary antibodies used in this study are shown in **Table S9**.

## Availability of data and materials

All raw sequence data are also available in the Genome Sequence Archive in the BIG Data Center, Beijing Institute of Genomics (BIG), Chinese Academy of Sciences, under accession number PRJCA004294. We also constructed the dog hippocampus atlas website at https://ngdc.cncb.ac.cn/idog/ (Single cell module).

## Declaration of Interests

The authors declare no competing interests.

## Author contributions

G.-D. W., S. Z. and B. M. conceived of the study, Q.-J. Z., X. L. and R. W. analyzed the data, Q.-J. Z. and L. Z collected the data, L. Z. performed experiment, X. L., G. L., Y. H., Z. D. and P. M. performed scRNA-seq library, T. Y helped anatomy experiment, S.-Z. W revised the manuscript. All authors wrote, revised, and approved the manuscript.

## Acknowledgement

This work was supported by the National Key R&D Program of China (2019YFA0707101, 2019YFA0709501), Key Research Program of Frontier Sciences of the CAS (ZDBS-LY-SM011, QYZDB-SSW-SYS008), National Natural Science Foundation of China (61621003), National Ten Thousand Talent Program for Young Top-notch Talents, and Innovative Research Team (in Science and Technology) of Yunnan Province (202005AE160012).G.D.W. is supported by the Youth Innovation Promotion Association of CAS. This work was supported by the Animal Branch of the Germplasm Bank of Wild Species, Chinese Academy of Sciences (the Large Research Infrastructure Funding).

## Supplemental Information

**Table S1 DEG list**.

**Table S2 GO enrichment analysis**.

**Table S3 630 PSGs expression in each cell type**.

**Table S4 Putative PSGs and DEGs statistical analysis**.

**Table S5 Interaction by GENIE3**

**Table S6 GO enrichment analysis result for regulatory genes**

**Table S7 Three differential expression analyses**.

**Table S8 Domesticated gene list**.

**Table S9 Antibody used in IF and IHC of Canis Hippocampus**.

**Table S10 Gene expression average and proportion**.

**Figure S1 Cluster analysis detail information**.

**Figure S2 The gene expression pattern of oligodendrocyte and molecular diversity between dog and mouse**.

**Figure S3 The mean expression of 630 putative PSGs in different cell types**.

**Figure S4 High-definition immunohistochemical staining**.

## Notes

### Competing Interest Statement

The authors have declared no competing interest.

