## supplementary for "Single-nuclei transcriptomics of dog hippocampus reveals the distinct cellular mechanism of domestication": supplementary figure.docx

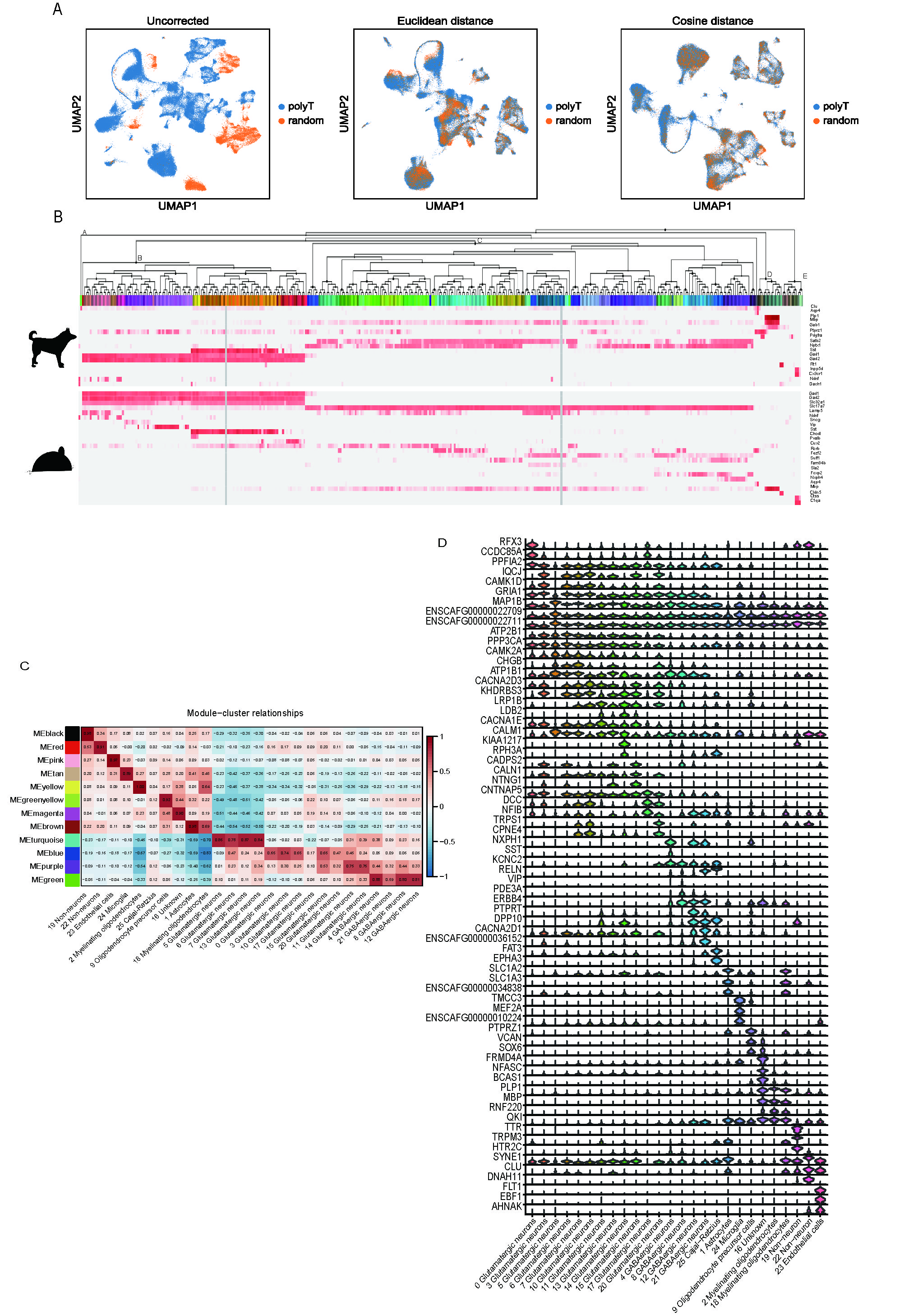

**Figure S1.** **Cluster analysis detail information.** (A) The echnical effect was corrected by partial PCA and cosine distance. (B) Dog cell types marker genes expressed in mouse brain atlas. Here are 17 genes as dog mark genes in our dog hippocampus data, including astrocytes (*Clu*, *Aqp4*), myelinating oligodendrocytes (*Plp1*, *Mbp*, *Gab1*), oligodendrocyte precursor cells (*Ptprz1*, *Pdgfra*), glutamatergic neurons (*Satb2*, *Nptx1*), GABAergic neurons (*Sst*, *Gad1*, *Gad2*), microglia (*Inpp5d*, *Cx3cr1*), and Cajal-Retzius cells (*Ndnf*, *Dach1*). Figure B also showed 24 mouse different cell types marker genes. Cluster A is Cajal-Retzius cells, cluster B is GABAergic neurons, cluster C is the glutamatergic neurons, cluster D is astrocytes and oligodendrocytes, cluster E is immune and vasculature (© 2015 Allen Institute for Brain Science. Allen Cell Types Database. Available from: <https://celltypes.brain-map.org/rnaseq/mouse_ctx-hip_10x>). (C) Gene modules detected by WGCNA and their enrichment with different cell types. (D) Vlnplot to show the DEGs expression profiles of each cluster.

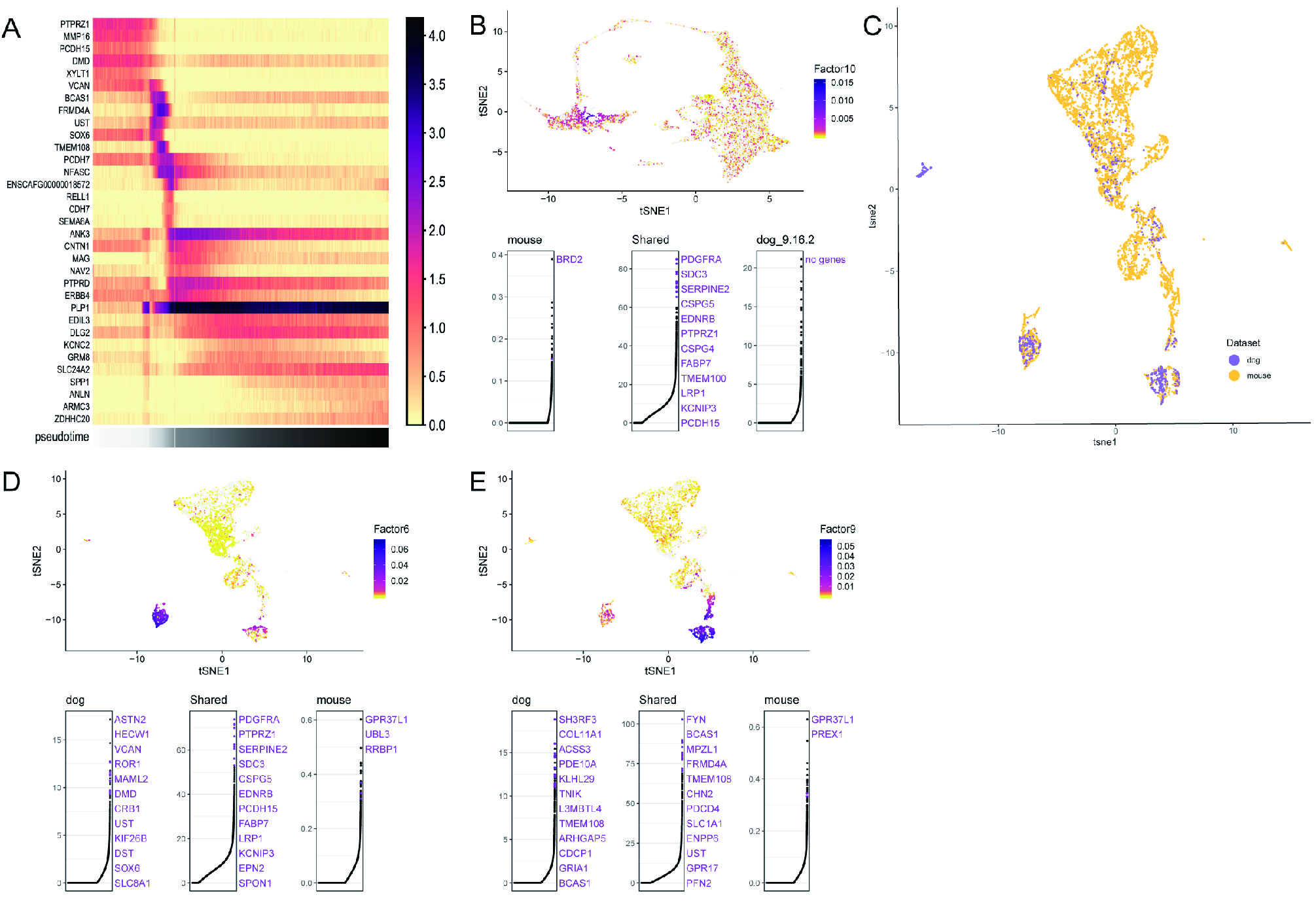

**Figure S2. The gene expression pattern of oligodendrocyte and molecular diversity between dog and mouse.** (A) Heatmap showing significantly expressed genes are sorted by pseudotime clusters sequence at the bottom. (B) Cell factor loading values (top) and gene loading plots (bottom) show dataset-specific and shared genes for factor 10. In gene loading plots, gene names are sorted in decreasing order of magnitude of their factor loading contribution and correspond to colored points in scatterplots. Plots are organized to show the metagene specific to the dog and mouse and the shared metagenes (PDGFRA, PTPRZ1) common to two datasets. (C) t-SNE visualization of 6,328 single cells (1,259 from dog and 5,069 from mouse) analyzed by LIGER, color-coded by species. (D–E) UMAP plots showing cell factor loading values (top) and gene loading plots (bottom) for factors corresponding to OPCs and differentiation-committed oligodendrocyte precursors (COPs). (D) Cell factor loading values (top) and gene loading plots (bottom) show dataset-specific and shared genes for factor 6. (E) Cell factor loading values (top) and gene loading plots (bottom) show dataset-specific and shared genes for factor 9. (B-E) Two OPC marker genes were found (*PDGFRA, PTPRZ1*) in both dog and mouse (**Figure S2B**). Since the identity of cluster 16 was unknown, we integrated cluster 16 with mouse cells only for further analysis (**Figure S2C**). As result, cluster 16 is defined as an intermediate state between mature cells and precursor cells. Specific component factors showed that *PDGFRA* and *FYN* shared in precursor cells (**Figure S2D and E**). However, they are two different precursor cell types’ markers in mice, which are OPCs and differentiation-committed oligodendrocyte precursors (COPs), and COPs lacked *PDGFRA* and *CSPG4* (Marques, et al. 2016).

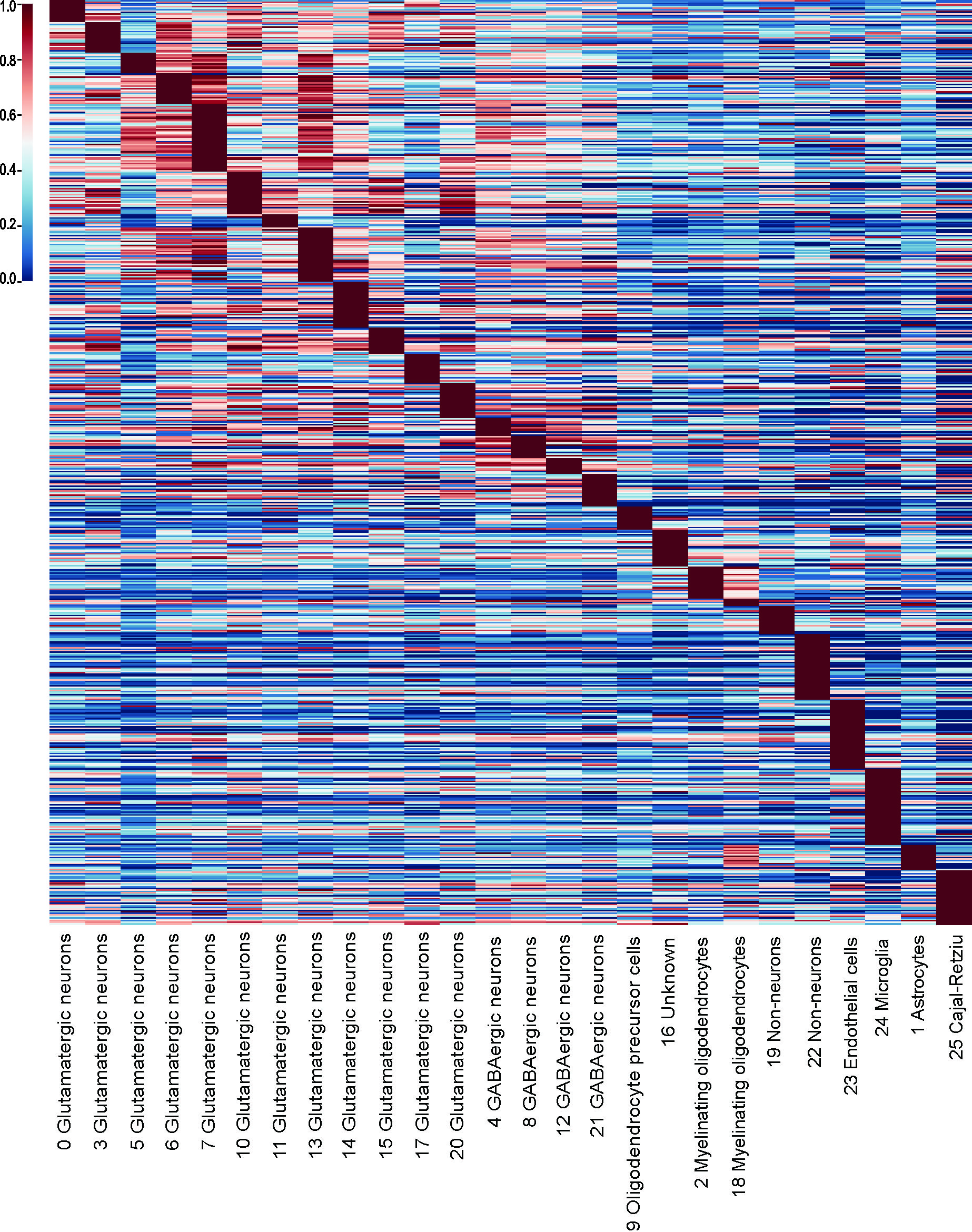

**Figure S3. The mean expression of 630 putative PSGs in different cell types.** The values of each gene (row) were its original ones divided by its maximum. This figure related to **Table S3.**

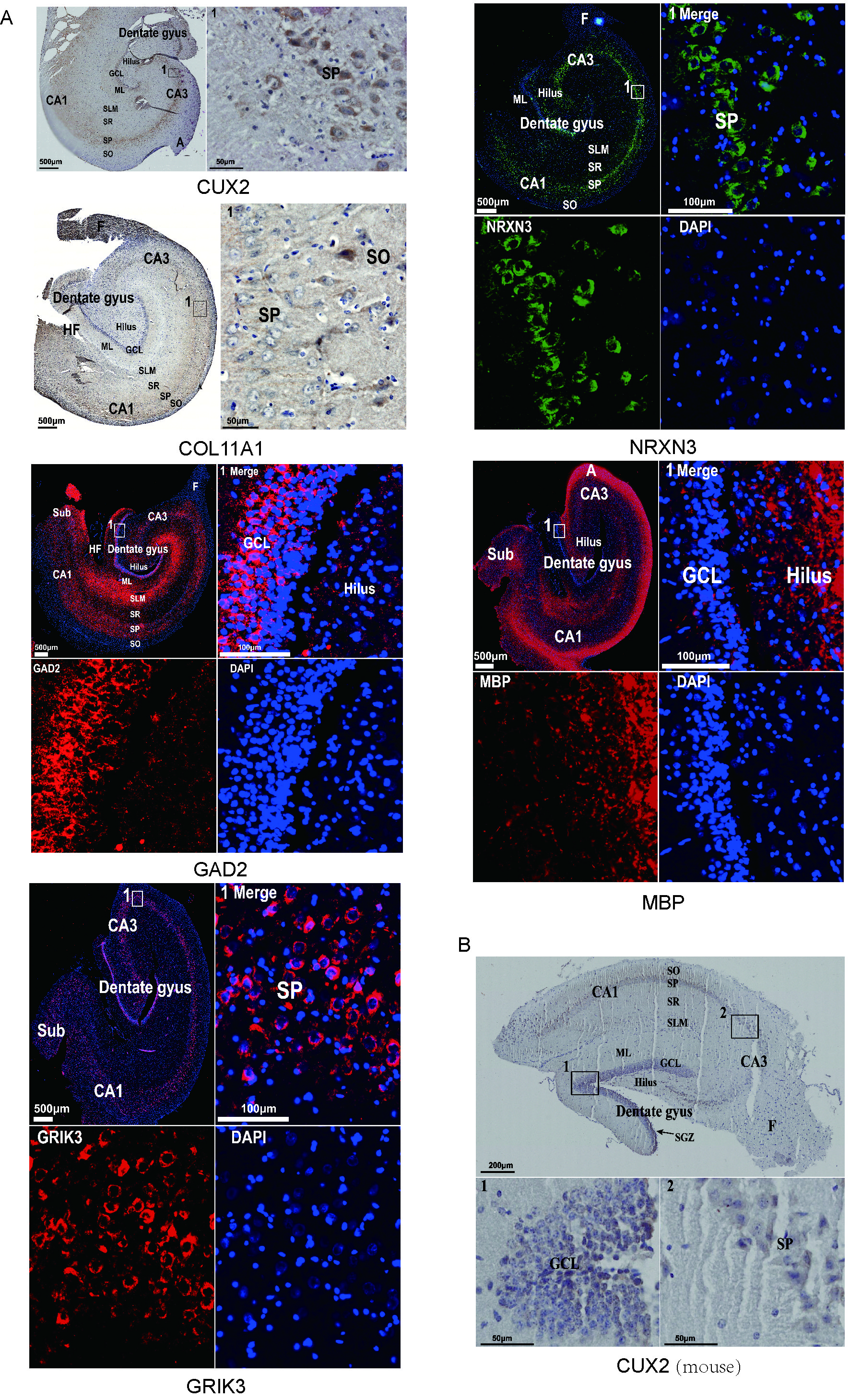

**Figure S4. High-definition immunohistochemical staining** (this figure is related to **Figure 3)**.

(A) Immunohistochemical staining: *CUX2*, *COL11A1*, *GAD2*, *GRIK3*, *NRXN3*, *MBP*.

(B) Visualization of CUX2 expression in mouse hippocampus by immunohistochemical staining. The expression of *Cux2* in the mouse hippocampus is the strongest in the subgranular zone (progenitor cell-enriched in this area) and hilus of DG cells, it is weakly expressed in CA3 and CA1 granular cells.
